# Evidence for multiscale multiplexed representation of visual features in EEG

**DOI:** 10.1101/2023.07.09.548296

**Authors:** Hamid Karimi-Rouzbahani

## Abstract

Distinct neural processes are often encoded across distinct time scales of neural activations. However, it has remained unclear if this multiscale coding strategy is also implemented for separate features of the same process. One difficulty is that the conventional methods of time scale analysis provide imperfect estimations of time scales when several components are active during a single process. Developing a novel decoding-based time scale estimation method, we detected distinct time scales for simultaneously present features of visual stimuli in electroencephalography. We observed that orientation and colour of grating stimuli were encoded in shorter whereas the spatial frequency and contrast of those stimuli were encoded in longer time scales. The conventional autocorrelation-based estimation of time scale was unable to detect these distinguishable time scales. These results provide new evidence for a multiscale multiplexed neural code in the human visual system and introduces a flexible method for estimating neural time scales.

## Introduction

Optimal decoding of neural codes remains an open question. This is because of the highly complex, dynamical, time-variant and often multiplexed patterns of the neural codes. Neural codes have been partially decoded in both invasive and non-invasive recording modalities. In invasive recordings, aspects of the neural spiking such as its rate, latency, temporal pattern or a combination of these have been proposed to represent the information about different cognitive processes (Borst and Theunissen, 1999; Panzeri et al., 2010). While rate coding proposes that information is transferred through the number of spikes within the encoding period (Theunissen and Miller, 1995; Denève and Machens, 2016; Merrikhi et al., 2021), latency coding conveys information through the relative latency of the spikes across multiple neurons (Gawne et al., 1996; Chase and Young, 2007) and temporal pattern coding reflects the information in precise timing of spikes within the encoding period (Mainen and Sejnowski, 1995; Butts et al., 2007).

While the literature has provided valuable insights about potential neural coding strategies in the brain, as they mainly used animal models (Victor, 2000; Kayser et al., 2009; Walker et al., 2011; Harvey et al., 2013), the generalisation of those insights to the human brain needs further research using non-invasive modalities. In humans, non-invasive recording modalities of electroencephalography (EEG) and magnetoencephalography (MEG) have provided excellent temporal resolution revealing the dynamics of information processing in the brain (Thorpe et al., 1996). Classical studies of EEG/MEG have used three sets of features of neural activations to access the neural codes from scalp-level brain activations. These include variance-based, frequency-based and information-theory-based features (Waschke et al., 2021), which comprise of a large array of features such as signal phase (Rupp et al., 2017), power across frequency bands as in time-frequency analyses (Majima et al., 2014), signal entropy (Richman and Moorman, 2000), complexity (Szczepański et al., 2003), etc. See Karimi-Rouzbahani et al. 2021a and Karimi-Rouzbahani et al. 2022, for systematic evaluation and comparison of these features in information coding.

With the development of machine-learning algorithms, there has been a shift from direct comparison of signal features (e.g., amplitude and power) across conditions of interest in individual channels (univariate), to quantifying their separability and generalisation to out-of-sample data across multiple channels (multivariate; Hebart and Baker, 2018). This neural decoding procedure, often called multivariate pattern decoding (MVPD), is generally performed over time in EEG/MEG to provide a temporally fine-grained view into the temporal dynamics of information coding in the brain (Grootswagers et al., 2017). To check the temporal dynamics of information coding across two conditions of interest e.g., orientation of two grating patches presented on the screen on two different sets of trials, one performs MVPD on every time point over the time course of the trials. The result will be a temporally changing information coding curve reflecting how distinctively and on which time windows the two conditions become discriminable in brain activations.

Despite the valuable insights that the traditional univariate and the more recent MVPD studies have provided, they generally make the implicit assumption that the neural processes of interest are encoded in similar time scales. Specifically, they assume that in order to utilise the high temporal resolution of EEG/MEG one needs to analyse every recorded sample across the trial separately and that would uncover the whole neural code. However, distinct cognitive processes have been shown to be encoded by the brain over different time scales. For example, we know from previous studies that while sensory information is generally processed along shorter time scales, information such as memory, reward, decision and motor are encoded along longer time scales (He, 2014; Cavanagh et al., 2020; Golesorkhi et al., 2021b; Soltani et al., 2021). Moreover, there is a correspondence between the time scales of neural processes and the dominant time scales of information processing across brain areas. Specifically, it has also been shown that dominantly sensory areas of the brain process information along shorter time scales whereas dominantly higher-order associative areas process information along longer time scales (He, 2014; Murray et al., 2014; Golesorkhi et al., 2021a; Soltani et al., 2021; Pinto et al., 2022; Li and Wang, 2022). This suggests that information from different processes and different brain areas do not necessarily appear within similar time scales.

We believe that distinct time scales do not only appear across distinct processes, but can also be generalised to distinct features of the same process such as different features of the same sensory input. This is supported by previous animal studies showing that stimulus contrast was represented at a temporal scale of about 10 ms, while stimulus orientation and the spatial frequency were encoded at a coarser temporal scale (30 ms and 100 ms, respectively; Victor, 2000). Moreover, spike rates on 5 ms to 10 ms timescales carried complementary information to the phase of firing relative to low-frequency (1–8 Hz) local field potentials (LFPs) about epoch of naturalistic movie (Montemurro et al., 2008). Despite differences between information encoding in animals and humans and the complex correspondences between invasive and non-invasive recordings (Ng et al., 2013; Panzeri et al., 2015), potential differences in the time scales of different conditions should be noted when decoding neural information in EEG/MEG. The implementation of multiscale information coding strategies can, among others, help the brain to multiplex several features of the process into a combined activity pattern leading to energy saving, robustness against noise and a higher capacity for information coding (Panzeri et al., 2010).

Here, we check to see if the human brain implements a multi-scale coding strategy to encode distinct features of sensory visual information at distinct temporal scales. To this end, we decoded four orthogonal aspects of information in a dataset where visual grating patches in different orientations, spatial frequencies, colours and contrasts were presented to participants in a passive-viewing rapid serial visual presentation paradigm in EEG. In addition to using the conventional approach of using the autocorrelation function (ACF) to quantify time scales of signals, we developed a novel decoding-based procedure which utilises the strength of MVPD to quantify the information within different time scales of simultaneously presented features of visual stimulus. We show that while the orientation and colour of the gratings were encoded in shorter time scales, the frequency and contrast of those gratings were encoded in longer time scales suggesting a multiplexed multiscale neural coding strategy in the human brain. To the best of our knowledge, this work provides the first human EEG evidence supporting previous animal studies which reported multiplex encoding of distinct stimulus features in sensory cortices (Victor, 2000; Kayser et al., 2009; Walker et al., 2011; Harvey et al., 2013). We also show that the conventional autocorrelation function is unable to discriminate the time scales of such entangled aspects of stimulus information. This is because rather than one time scale per feature affecting the recorded time series, the ACF-based methods detect a combined time scale for all features simultaneously present in the trial.

## Results

Using multivariate pattern decoding, we looked for multiscale neural codes in four simultaneously present orthogonal features of visual stimuli (Figure 1A). We evaluated 10 time scales ranging from 2ms to 74ms to see if those features were encoded at different time scales. EEG was collected while visual grating stimuli were presented to participants in a passive rapid serial visual presentation paradigm (Figure 1B; dataset from Grootswagers et al., 2023a). We performed pairwise decoding of four orientations, four spatial frequencies, four colours and four contrast levels over time. Each stimulus presented one of 256 combinations of the four features and the decoding of each feature was orthogonal to other features. To check the level of information in different time scales, we temporally averaged the signal time series using moving average windows of one of 10 different lengths/time scales (Figure 1C). We also tested the more conventional method of time scale estimation based on signal autocorrelation.

**Figure 1.**
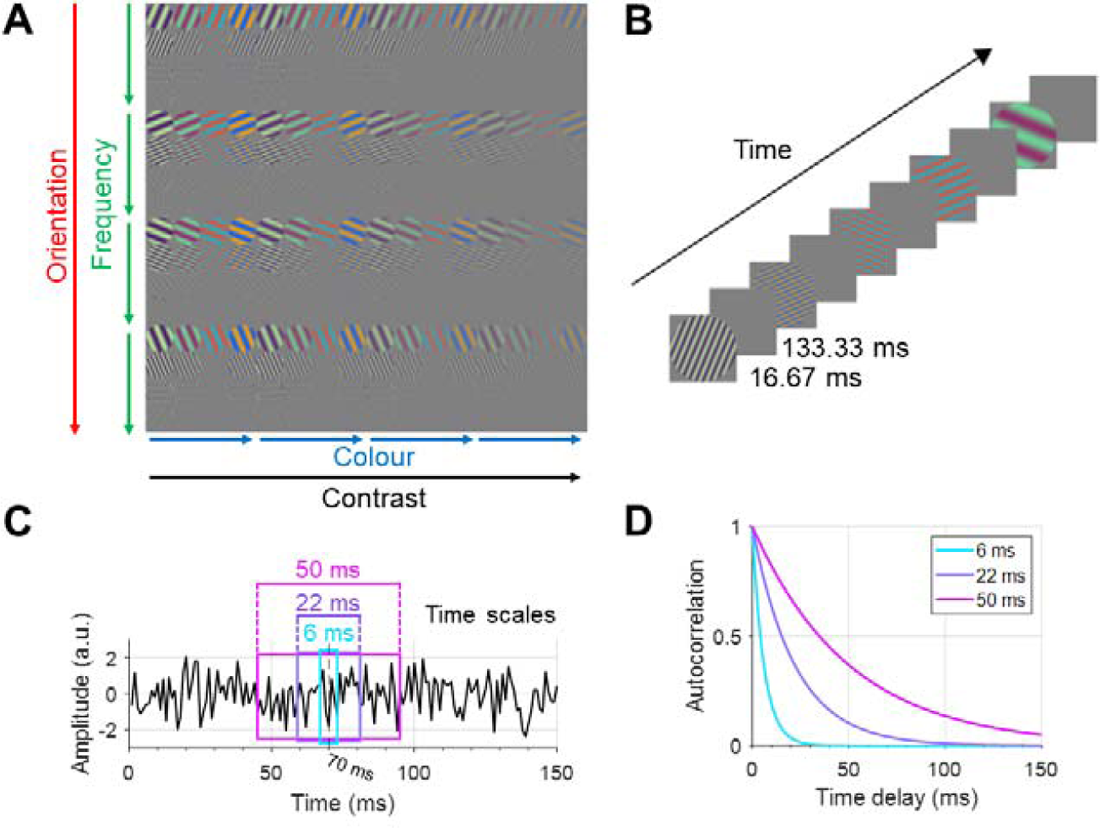
Stimulus set, experimental paradigm and time scale estimation methods. A and B modified with permission from Grootswagers et al., 2023b. (A) 256 grating stimuli were generated each with one of four conditions of the features of orientation, colour, spatial frequency and contrast. (B) a rapid serial visual presentation paradigm was used in the EEG recording, where each stimulus was presented for 16.67ms followed by 133.33ms inter-stimulus interval. (C) a simulated time series showing the lengths of three of our evaluated time scales of 6ms, 22ms and 50ms centred on the time sample of 70ms. (D) simulated exponentially decaying functions (^_^) which reflect typical autocorrelation functions corresponding to time scales of 6ms, 22ms and 50ms (time scale = the time delay at which the autocorrelation function is 0.368 times its initial value).

### Temporal dynamics of information coding across time scales

For clearer presentation of time resolved decoding results, we only present the results for time scales of 6ms (short), 22ms (medium) and 50ms (long) in the main text. For other time scales see Supplementary Figure 1.

There was evidence (Bayes Factor; BF > 3) for above-baseline decoding for all four features after around 80ms post-stimulus onset for all time scales, which remained above baseline until around 300ms for orientation, 380ms for colour and until 500ms for both frequency and contrast information (Figure 2). Generally, evidence for above-baseline decoding appeared earlier for longer vs. shorter time scales (see BFs comparing 22ms and 50ms time scales in Figure 2 and against baseline in Supplementary Figure 1).

**Figure 2.**
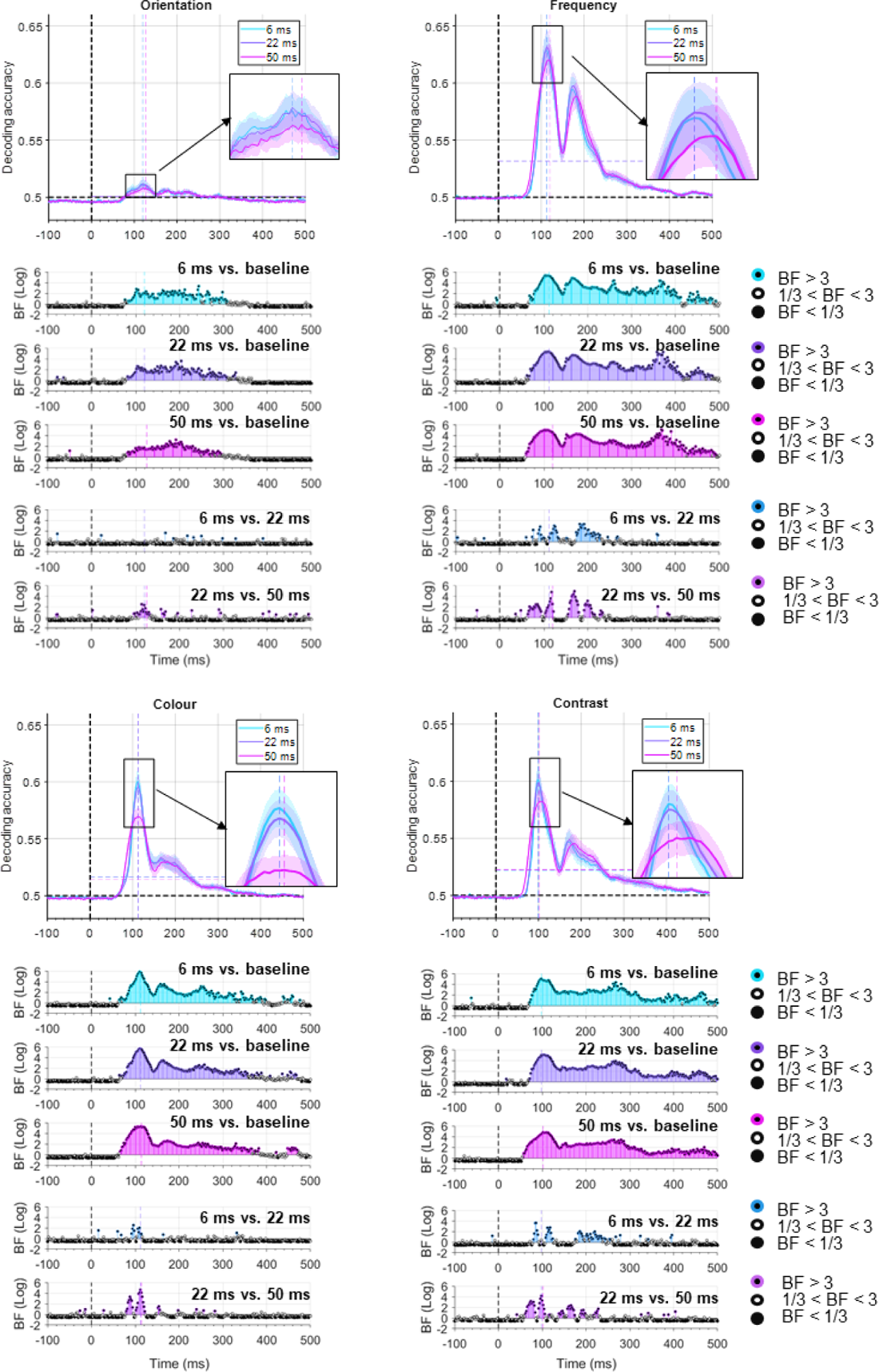
Decoding of stimulus features in three representative time scales. Decoding time courses are shown for orientation (top left), colour (bottom left), frequency (top right) and contrast (bottom right). Top panels: solid decoding lines and shadings reflect average and standard error across subjects, respectively. Black horizontal dashed line indicates theoretical chance-level decoding (0.5). Black vertical dashed line indicates stimulus onset time. The vertical and horizontal coloured dashed lines indicate the time of maximum decoding and the average post-stimulus decoding for the corresponding time scales (note that they may overlap), respectively. Bottom panels: Bayesian evidence for difference from baseline (i.e., −100 to 0ms) and difference between time scales. Bayes factors calculated using Bayes t-test (BF): Coloured filled circles show evidence for difference (BF > 3) or against any difference (BF < 1/3) and empty circles indicate insufficient evidence (1/3 < BF < 3). Note that colours were averaged when

The first peak of decoding occurred within in the 90ms-130ms window as indicated by the coloured vertical dashed lines (note that some overlap) and enlarged panels in Figure 2. At decoding peaks, we observed the following. In orientation decoding, there was either evidence against (BF < 1/3) or insufficient evidence (1/3 < BF < 3) for higher decoding of orientation in 6ms than 22ms time scale or vice versa. In colour decoding, there was evidence (BF > 3) for higher decoding in 6ms than 22ms time scale. In frequency decoding, there was evidence (BF > 3) for higher decoding in 22ms than 6ms time scale. In contrast decoding, there was evidence against (BF < 1/3) higher decoding in 6ms than 22ms time scale or vice versa. In all features, there was evidence (BF > 3) for higher decoding in 22ms than 50ms time scale.

After the first peak, we made these observations. In orientation and colour decoding, there was no cluster of time points with evidence (BF > 3) for higher decoding in one of the three time scales compared to the others. In frequency decoding, there was evidence (BF > 3) for higher decoding in 22ms than 50ms time scale before (160ms-180ms) and higher decoding in the 6ms time scale after the second peak (180ms-240ms). There was evidence (BF > 3) for higher decoding in 50ms than 22ms time scale after the second peak (180ms-240ms). In contrast decoding, the same pattern was observed with evidence (BF > 3) for higher decoding in 22ms than 50ms around the second peak (160ms-180ms) followed by evidence (BF > 3) for higher decoding in 50ms than 22ms and in 22ms than in 6ms time scale in the following windows (180ms-250ms). These results show that not all features provide their maximal information in similar time scales, with orientation showing no clear difference between short (i.e., 6ms) and medium (i.e., 22ms) time scale, colour dominantly encoded in short (i.e., 6ms), and frequency and contrast dominantly encoded in medium (i.e., 22ms) time scale across time. Condition-specific decoding curves also showed the same pattern of time scale preference (Supplementary Figure 2).

### Dominant time scales of neural code for each feature

The results presented above suggested that there might be a dominant time scale for each of the four features supporting the idea of a multiplexed multiscale coding of information. To check that, we extracted the peaks and averages of decoding curves as two summary statistics from the decoding curves presented in Figure 2, normalised them within each feature across time scales and overlayed them in the same panel to facilitate their comparison. We incorporated the original 10 time scales for a fine-grained view. This gave us a time scale tuning-curve, which resembled those obtained from single/multi-neuron data used to check neural selectivity (Figure 3; Merrikhi et al., 2021). The reason for using both peak and average of decoding is that, it is not yet clear which aspect of a time-resolved decoding curve reflects the true nature of the neural code and its dynamics (Grootswagers et al., 2017; Karimi-Rouzbahani et al., 2017b; Karimi-Rouzbahani et al., 2018; Karimi-Rouzbahani et al., 2021a). Therefore, we explain both statistics in Figure 3. Time scale tuning curves showed inverse U-shaped time scale curves for different features supporting that there are “preferred/dominant” time scales for each feature (Figure 3A).

**Figure 3.**
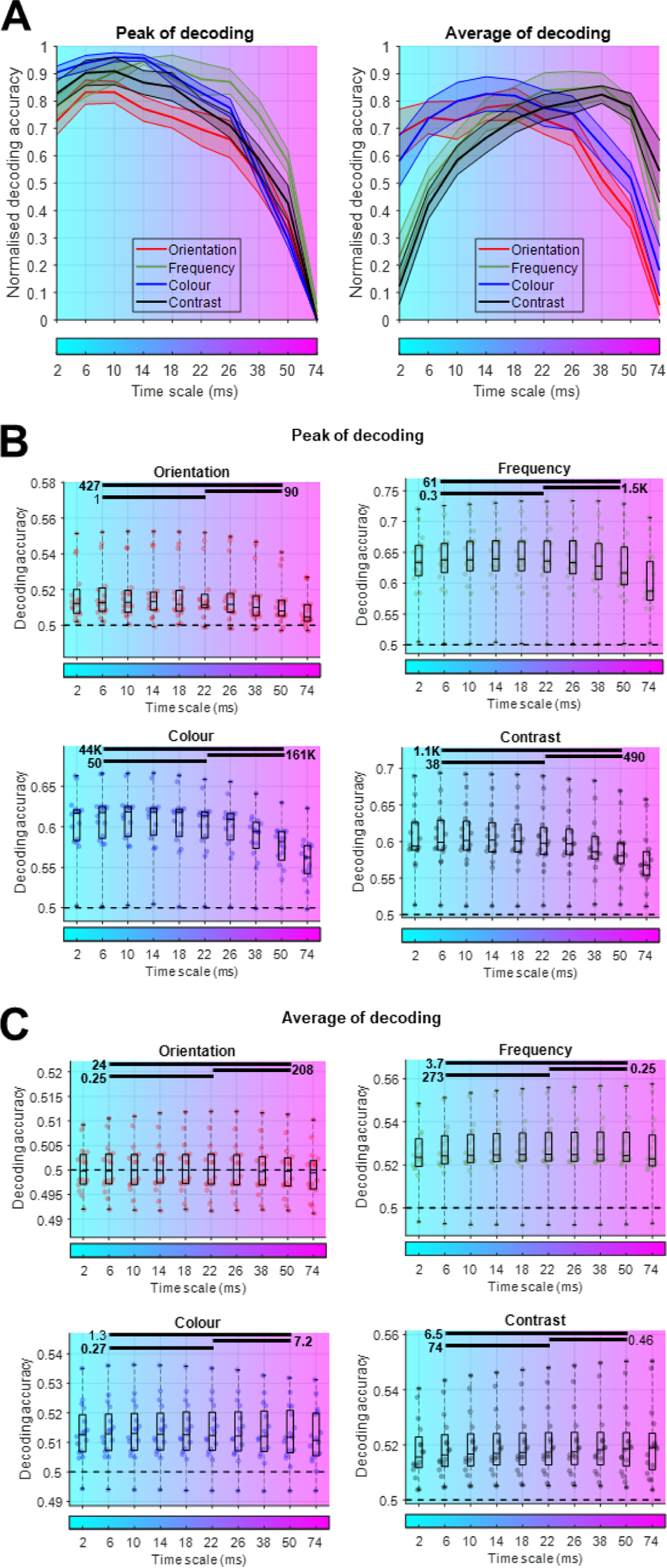
Comparison of summary statistics of decoding curves across time scales and features. (A) normalised peak and average decoding for each of the four features across 10 time scales. For average decoding, the data in the post-stimulus span was averaged. Solid decoding lines and shadings reflect average and standard error across subjects, respectively. Normalisation of decoding was performed over time scales within each subject. Time scales are shown using a spectrum of colours along the horizontal axis which matches the time scale colours in previous figures. (B) original (un-normalised) peak of decoding accuracies across time scales. Box plots show the distribution of data, its quartiles and median and whiskers indicate the maximum and minimum of the data over subjects. Each dot indicates the data from one subject. Horizontal dashed line refers to theoretical chance-level decoding (0.5). (C) same as (B) for average of decoding. Solid horizontal lines indicate the time scales which are compared (i.e., 6ms, 22ms and 50ms) and numbers indicate Bayesian evidence for the difference between those time scales. Evidence for and against difference are marked using bold fonts.

Time scale tuning curves based on peaks of decoding showed a longer dominant time scale for frequency than the other features (Figure 3A, left). Interestingly, the differences between features become clearer when looking at the tuning curves obtained from average of decoding (Figure 3A, right). While orientation and colour showed a shorter dominant time scale (18ms and 14ms, respectively), frequency and contrast showed longer dominant time scales (both around 38ms). We also present the original un-normalised peaks and averages of decoding data and their Bayesian statsitical evaluation (Figures 3B and 3C). Here we sufficed to statistically compare the representative time scales of 6ms, 22ms and 50ms to show the general trend. In peak of decoding data (Figure 3B), while there was evidence (BF > 38) for a drop from 6ms to 22ms for colour and contrast features, there was insufficient evidence (BF = 1) or evidence (BF = 0.3) against difference between those time scales in orientation and frequency, respectively. In average of decoding data (Figure 3C), there was distinctive effects of time scales across features. There was evidence (BF < 0.27) against any change in decoding between 6ms and 22ms for orientation and colour, and evidence (BF > 7.2) for decrease in decoding from 22ms to 50ms for both features. On the other hand, there was evidence (BF > 74) for an increase in decoding from 6ms to 22ms for both frequency and contrast and evidence (BF = 0.25) against or insufficient evidence (BF = 0.46) either way for any change in decoding from 22ms to 50ms time scales in frequency and contrast decoding, respectively. These repeat the results in Figure 3A: while orientation and colour were dominantly encoded in shorter time scales, frequency and contrast were dominantly encoded in longer time scales. These results support a multiplexed multiscale strategy implemented by the brain to encode some features in time scales different from the others.

### Multiscale encoding of information across time

The results shown above suggest that each feature (e.g., frequency) might be decodable in different time scales depending on how far into the trial the analysis is performed. Specifically, we observed that around the first peak (∼180ms), frequency decoding was maximal in the 22ms time scale, but after the second peak it was maximal in the 50ms time scale. This makes us wonder if each feature is dynamically encoded across multiple time scales over the course of the trial. This can be supported by the fact that different neural mechanisms can get involved in the processing of visual stimulus each with different time scales (Panzeri et al., 2012; Golesorkhi et al., 2021). The other question is, despite each feature having its own dominant time scale across the trial, whether there is any similarity between the temporal dynamics of dominant time scales across features - are there general feature-independent mechanisms supporting multiscale coding across features? To shed light on these questions, we calculated a multiscale decoding curve for which we only retained the maximum decoding accuracy from the best-performing time scale on every time point along the trial (Figure 4) and reported the corresponding time scale across time. Results showed that, rather than one individual time scale, a wide range of time scales contributed to the multiscale decoding curve which dynamically changed over the course of the trial. Across features, there were similarities between the timing of best-performing time scales. Specifically, longer time scales were dominant during the initial ramp up of the multiscale decoding curves followed by the dominance of short temporal scales at the peak across all features (Figure 4). This might reflect a common dynamical encoding mechanism across features which encodes sensory inputs as they arrive with different delays from the eye over longer time scales (e.g., different contrasts arriving with different delays (Oram, 2010; Zamarashkina et al., 2020; Himberger et al., 2018)) followed by fine-grained information encoding in shorter time scales when majority of sensory inputs are present. Despite these similarities, the multiscale decoding results were not identical across features. In accordance with the results in Figure 3, while multiscale decoding of orientation and colour features were more frequently supported by shorter time scales (< 22ms), frequency and contrast features were more frequently supported by longer time scales (> 22ms; Figure 4 and Supplementary Figure 3). This supports our earlier results regarding distinct dominant time scales supporting the encoding of different features (Figure 3).

**Figure 4.**
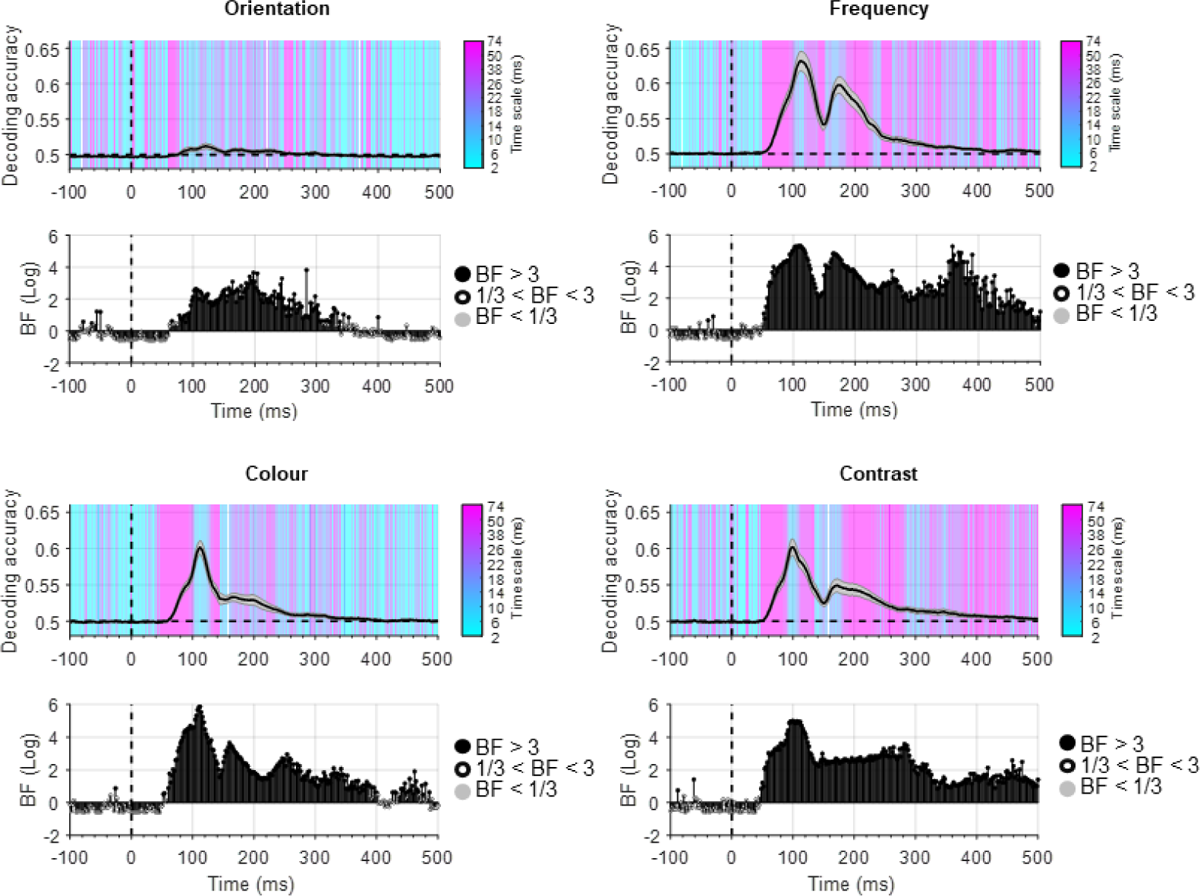
Multiscale decoding of information. Decoding time courses are shown for orientation (top left), colour (bottom left), frequency (top right) and contrast (bottom right). Top panels: solid decoding lines and shadings reflect average and standard error across subjects, respectively. Horizontal dashed line refers to theoretical chance-level decoding (0.5). Black vertical dashed lines indicate stimulus onset time. A spectrum of colours indicates the time scale which provided the maximum decoding performance on every time point, which is used to generate the combined multiscale decoding curve. Bottom panels: Bayesian evidence for difference from baseline (−100 to 0ms). Bayes factors calculated using Bayes t-test (BF): filled circles show evidence for difference (BF > 3; black) or against any difference (BF < 1/3; grey) and empty circles indicate insufficient evidence either way (1/3 < BF < 3).

### Univariate estimation of time scales

Using a novel multivariate pattern decoding method, we showed that distinct features could be encoded at distinct time scales. As a final step, we checked to see whether we could have detected the time scales of these features using a more conventional approach for estimating time scales - the ACF-based estimation (Golesorki et al., 2021b; Zeraati et al., 2022). This conventional approach first calculates the ACF of the time series/neural activations, and then estimates its time scale by fitting an exponential decay function to it (see Methods). However, obviously this method would not be useful in this work as the aim here was to estimate the time scale for each feature separately, which were all present simultaneously within the same trial. Therefore, the estimated time scale would be similar for all features as their trials are identical. We estimated the time scales of every EEG channel and every trial using the conventional ACF-based estimation. This serves two purposes. First, it allows us to test one of the most stablished findings in time scale analysis in the brain which states that lower order sensory areas have a shorter time scale than higher-order association areas (Golesorkhi et al., 2021b). Second, it allows us to see if our decoding-based estimated time scales fall within the same range as the ACF-based estimations, validating our proposed method.

To see if the time scales increase from the sensory-dominated areas to higher-order areas in EEG, we analysed average time scales across three regions of interest (ROI) along the posterior to anterior scalp electrodes (Figure 5A). ROIs were selected in a way to avoid the very frontal electrodes which might be affected by residual eye-movement artefacts despite ICA-based artefact removal. While there was insufficient evidence (BF = 0.64) for any increase in time scale from occipital to central ROI, there was evidence for longer time scales in frontal than in central (BF = 15.6) and occipital (BF = 17.2) ROIs. This supports previous neural data (Soltani et al., 2021; Pinto et al., 2022; Li and Wang, 2022) and recent MEG/EEG (Golesorkhi et al., 2021a) studies showing that neural time scales increase from posterior sensory to frontal association areas.

**Figure 5.**
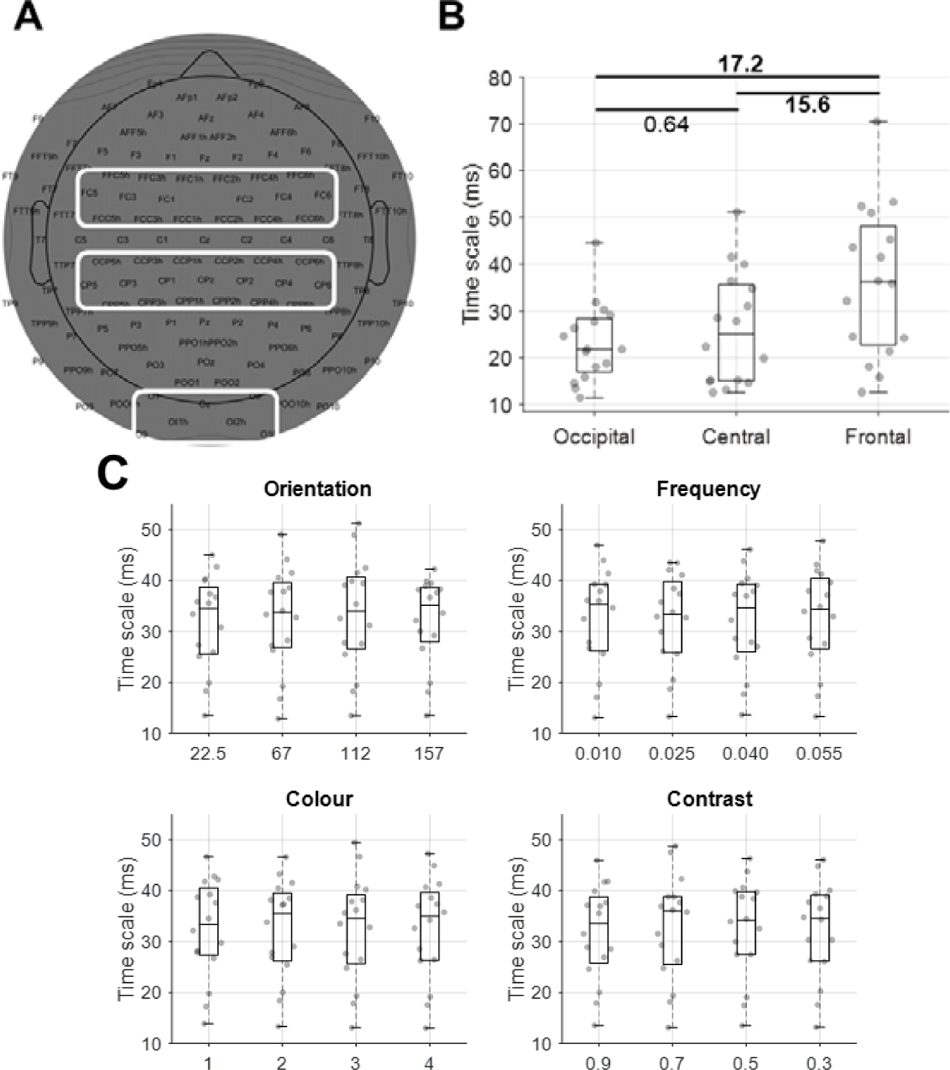
Estimation of time scales using autocorrelation function. (A) scalp electrode positions, selected as occipital, central and frontal ROI. (B) estimated time scales obtained by averaging the time scales within the three ROIs shown in (A). Box plots show the distribution of data, its quartiles, median and whiskers indicate the maximum and minimum of the data over subjects. Each dot indicates the data from one subject. Numbers indicate Bayesian evidence for the difference between ROIs (B). Evidence for and against difference are marked using bold fonts. (C) shows estimated time scales from the whole head for each feature and its four conditions. There was no evidence (BF < 3) for any difference between conditions in any of the four features.

Next, we averaged the estimated time scales from all electrodes separated by conditions within each feature (Figure 5C). We observed that the electrode-averaged estimated time scales ranged from 12ms to 52ms across conditions and features overlapping with the range which obtained maximal average decoding in our proposed decoding-based time scale estimation method (Figure 3A). This validates our method and highlights its flexibility in estimating time scales from simultaneously active processes. We also statistically checked to see there was a difference between the estimated time scales for conditions of each feature, but there was either no (BF < 1/3) or insufficient evidence (1/3 < BF < 3) for any difference between conditions. This is because every condition in each trial is accompanied by other conditions from other features which moves the estimated time scale towards a cross-condition combined time scale.

## Discussion

This study developed a new method to gain insights into multiscale neural coding mechanisms of the human brain in sensory visual processing as reflected in EEG. This work makes three major contributions to the investigation of information coding the human brain, which we discuss below and compare with the relevant literature.

First, we show that not all features of low-level grating stimuli were encoded in similar time scales. While previous studies have established that distinct neural processes can have different time scales (He, 2014; Cavanagh et al., 2020; Golesorkhi et al., 2021a), it has remained unknown if it would generalise to the components of a single process (e.g., sensory processing). Multiplexed multiscale neural codes have been observed in neural data from ferret (Walker et al., 2011) auditory cortex as well as monkey (Kayser et al., 2009) auditory, visual (Victor, 2000) and somatosensory (Harvey et al., 2013; Rossi-Pool et al., 2021) cortices. However, to the best of our knowledge, such multiscale coding of information has not been reported for visual sensory coding in human EEG. In fact, this makes our results even more important because a direct correspondence between invasive neural recording and non-invasive EEG is unclear. For example, it has been shown that neural firing rates in invasive recording reflect in the phase patterns rather than the intuitive amplitude of EEG oscillations (Ng et al., 2013).

Our findings complement a few recent EEG decoding studies. One of them compared a large set of signal features, ranging from low-level signal moments to high-level phase and amplitude features obtained through Hilbert and Wavelet transforms, and showed that subtle sample-to-sample signal variabilities in longer time windows (50ms), which are often overlooked through down-sampling, provide additional information about semantic object categories than the down-sampled signal amplitudes (Karimi-Rouzbahani et al., 2021b). This added to earlier evidence suggesting that temporally variable patterns of activation can provide a parallel complementary platform for information coding along signal amplitude (Orbán et al., 2016; Karimi-Rouzbahani et al., 2017a). Our work complements these previous studies by showing that, informative temporal variabilities can encode information over different time scales. Our current results align with a recent study showing that multiscale patterns reflect the neural codes better and can predict the behaviour more closely than a combination of multiple distinct encoding schemes (Karimi-Rouzbahani et al., 2022). Although there has been evidence for more information in medium-scale (50ms) than shorter (10ms) or longer (100ms) temporal scales in encoding of object categories (see Supplementary Figure 3 in Karimi-Rouzbahani et al., 2021b), no studies have made systematic evaluation of multiscale encoding of simultaneously present features of visual stimuli in EEG.

Another contribution of this study is that it introduces a novel method for estimating time scales of neural processes. This method relies on activity within windows of different temporal scales and is especially important for situations, where conditions of interest (i.e., different visual features), are temporally adjacent, or such as in this work simultaneously present, making it difficult to evaluate the time scale of each individual condition separately. The proposed method directly computes the information contained within each time scale about each neuronal process and the features within that process. As we showed above, by comparing the amount of information within each temporally averaged time window of activation, we obtained insights about the most informative time scale of each feature. Conventionally, several methods have been used to estimate time scales of neuronal processes. The dominant method is based on fitting a decaying exponential function to the autocorrelation function (Golesorkhi et al., 2021b), which suffers when the length of the time series is short (Zeraati et al., 2022). We showed here that, autocorrelation-based estimation of time scales is unable to distinguish between our overlapping features as they all affect the neural activation simultaneously (Figure 5). Moreover, the estimation error in ACF-based estimation increases in proportion to the number of simultaneous processes as they all affect the time series causing a hard-to-track pattern in the AFC. Another method based on generative Bayesian frameworks has been proposed which allows the estimation of more than one time scale from an individual time series/neural activation (Zeraati et al., 2023). This method needs prior knowledge about plausible generative models of the underlying processes which might be hard to obtain and remains silent about the one-to-one correspondence between the estimated time scales and the underlying processes. The shape of the power spectrum has also been used to estimate the time scale of neural processes. However, this method is more suited for estimating time scales of oscillatory processes (Donoghue et al., 2020). Another time scale estimation method relies on fitting autoregressive models to the time series data, which in a recent work has been used to successfully estimate a range of short (within-trial intrinsic) to longer (across-trial seasonal) time scales (Spitmaan et al., 2020). The performance of this method in estimating time scales of overlapping processes needs further investigation.

Our method, on the other hand, does not need prior knowledge about potential time scales, does not depend on the length of the time series, and can estimate the time scales even if the underlying processes occur simultaneously (i.e., features of the same stimulus). It is of note that, our proposed method of temporal windowing followed by multivariate decoding, does not need to necessarily average the samples of the time series to estimate the time scales. We used averaging as one of the most intuitive ways to summarise the neural code, which also has biological plausibility (Himberger et al., 2018; Norman-Haignere et al., 2022; Wolff et al., 2022). One can summarise the samples within the window by extracting a wide range of quantifiers, such as signal complexity (e.g., entropy, fractal dimension) or even using the original samples (Karimi-Rouzbahani et al., 2021b). Moreover, our decoding-based method of time scale estimation does not need to necessarily incorporate more than one activity channel – one can apply it to individual signal activations to gain insights into the spatial organisation of time scales as well. Therefore, our proposed method, provides a flexible and intuitive approach for estimating time scales of neural processes.

The third contribution of this study is the insights it provides about the advantage of doing multiscale rather than uniscale analysis when the time scales of the underlying processes are unknown, which is often the case. Generally, a constant time scale is implicitly selected across all conditions under study (Grootswagers et al., 2017). However, we showed that for more optimal decoding of each condition, it is worth evaluating a range of potential time scales not to miss the true dynamics of the process at hand. Here, we believe that we showed one of the most extreme cases, where distinct features of the same process (i.e., encoding of grating stimulus), were encoded across distinct time scales. In the case of distinct processes (e.g., sensory vs. memory), the differences between dominant time scales can potentially be much larger and might have more pronounced effects on our interpretations if overlooked.

Why does the brain generate multiplexed multiscale neural codes? It has been suggested that a multiplexed multiscale encoding scheme can provide several computational advantages over a fixed uniform neural code. This includes increasing the capacity of the brain to encode information by transferring multiple types of information within the same packet of data (Schaefer et al., 2006; Luczak et al., 2015). Moreover, multiscale neural codes were shown to have higher robustness to environmental noise which often accompanies sensory inputs in real-life situations (Kayser et al., 2009). In addition, brain has been suggested to reduce the ambiguity of the input stimuli using multiple simultaneous coding strategies (Schaefer et al., 2006; Schroeder and Lakatos, 2009). Therefore, multiscale coding provides a platform for a more optimal and robust encoding of information in the brain.

What underlies the similarities between the temporal dynamics of optimal time scales across different features? We showed that, for all four stimulus features (i.e., orientation, frequency, etc.), longer time scales were more optimal during the ramp-up window of the decoding followed by shorter time scales around the peak (Figure 4). The initial longer time scales might allow for the encoding of distinct conditions of each feature arriving with different delays (Himberger et al., 2018). For example, previous studies have shown that higher-contrast stimuli lead to earlier responses in visual areas than lower contrast ones (Oram, 2010), which generalises to other features of the visual stimuli too (Zamarashkina et al., 2020; Mazer et al., 2002). This is generally referred to as the temporal receptive field, which defines the temporal window in which the neuron encodes the input stimuli (Hasson et al., 2015). The peak decoding, on the other hand, possibly reflects the time at which most conditions within each feature (e.g., different levels of contrast) reach the visual neurons and are encoded most effectively. These interpretations imply that, as different conditions of a feature lead to different neural latencies, to be able to make inferences about the temporal dynamics of a feature based on its decoding pattern, one needs to evaluate a large set of conditions within each feature/experimental condition.

One future step will be to evaluate not only the time scales which lead to the highest peak and average in decoding, but to check if they provide the best prediction of behavioural performance. In fact, an enhancement in decoding performance does not necessarily mean a better access to the task-relevant neural code. It can be that the decoded information is not used by the brain to undertake the neural process at hand. One way to validate the plausibility of neural codes is to check if they reflect the behavioural performance in the task (Ritchie et al., 2015; Karimi-Rouzbahani et al., 2021b). While the dataset used in this study did not allow us to evaluate the correspondence between the neural code and behaviour, datasets with behavioural components corresponding to the neural process will allow evaluating the correspondence between the neural code and behaviour (Karimi-Rouzbahani et al., 2019; Karimi-Rouzbahani et al., 2021b; Karimi-Rouzbahani et al., 2022).

The overall goal of this study was to check if simultaneously presented features of the visual inputs are encoded across multiple time scales providing evidence for multiplexed multiscale coding of information in the human brain in EEG. We developed an intuitive decoding-based method for estimating time scales in times series and showed that distinct features of the same grating visual stimulus were decoded in different time scales – spatial frequency and contrast were decoded more optimally in longer time scales than orientation and colour. These results provide new evidence for multiplexed multiscale coding of sensory information in humans. The proposed time scale estimation method can be used to systematically evaluate time scales of a wide range of neural processes.

## Methods

### Dataset and pre-processing

We used a previously collected EEG dataset (Grootswagers et al., 2023a). The details of the dataset can be found in the original publication (Grootswagers et al., 2023b). Briefly, sixteen participants (11 female, 5 male, 18-27 years) took part in a rapid serial visual presentation paradigm in which a set of 256 grating stimuli (Figure 1A) were flashed on the screen (6.5 × 6.5 degrees of visual angle) for 16.67ms followed by 133.33ms blank screen (Figure 1B). While the original study had 50ms and 150ms inter-stimulus interval (ISI) trials, we only used the latter to allow implementing the classical time scale estimation method using more time samples. During the visual presentations, EEG signals were recorded using a 128-channel BrainVision ActiChamp system. Gratings were in one of four orientations, one of four spatial frequencies, one of four colours and one of four contrasts (256 combinations). Signals were collected at 1000 Hz and referenced to FCz online.

Pre-processing steps used here are different from the original study. Pre-processing was done in EEGLAB (Delorme and Makeig, 2004) and involved notch-filtering (FIR filters) at 50, 100 and 150Hz, band-pass filtering in the range from 0.05 to 200 Hz (FIR filters) and down-sampling to 500Hz. ICA was applied to remove eye-artefacts using the runica method as implemented in EEGLAB. The selection of artefactual channels were done in EEGLAB using ICLabel plugin and 2 components were removed on average per participant. Signals were epoched from −100 to 500 ms around the stimulus onset time. The analysis scripts can be found at https://github.com/HamidKarimi-Rouzbahani/Multiscale-multiplexed-coding-of-visual-features-in-EEG.

### Multivariate pattern decoding

We used a standard multivariate pattern decoding to quantify the level of different types of information consisting of orientation, frequency, colour and contrast in EEG. We decoded all pairwise combinations of four conditions within each feature. For example, in orientation decoding we decoded the 1^st^ against 2^nd^, 1^st^ against 3^rd^, 1^st^ against 4^th^, 2^nd^ against 3^rd^, 2^nd^ against 4^th^ and 3^rd^ against 4^th^ orientation and averaged before reporting them as orientation decoding. The same procedure was followed for all other features. We used a Linear Discriminant Analysis (LDA) classifier and repeated the decoding analysis on every time point along the trial. We used a 10-fold cross-validation procedure, with 9 folds of the data used for training the classifier and the left-out fold used for testing it and repeating the procedure 10 times until all folds were used once in testing. The decoding accuracy for a single participant was calculated by averaging the decoding results across these cross-validation runs.

### Estimation of time scales using multivariate decoding

To obtain insights into the importance of time scales in information coding, before the decoding was performed on signals, the signal time series for each channel and trial was temporally averaged at different scales. Specifically, signals were averaged across time using a moving average sliding time window with 10 different lengths (i.e., time scales; Figure 1C). This included windows of size 2ms (only one sample/no temporal averaging), 6ms (3 samples), 10ms (5 samples), …, 74ms (37 samples). This range was selected based on preliminary results - shorter or longer averaging windows would reduce the decoding accuracy. This multiscale averaging procedure resulted in signals with more (in the case of 2ms window) or less (in the case of 74ms window) temporal variabilities for decoding. The rationale behind this averaging procedure is that, the informative pattern (i.e., neural code) can be best detected if its length matches the length of the averaging window. Specifically, if the neural code, which is reflected in variabilities of EEG time series, is relatively short e.g., 6ms, using a long averaging window (e.g., 50ms) will suppress it by mixing it with uninformative variabilities in the averaging process and lead to reduced decoding. Here, a shorter averaging window at the same length of the neural code (e.g., 6ms) would maximise the decoding. On the other hand, if the neural code is relatively long e.g., 50ms a short averaging window will miss the long neural code as it will only see a portion of it and will lead to declined decoding. Here, a long averaging window (e.g., 50ms) can cover the neural code and lead to improved descent decoding. Please note that the indicated time in the time-resolved decoding results reflects the middle sample in the window. Accordingly, decoding at time 70ms reflects the result of averaging samples 68ms, 70ms and 72ms in the case of 6ms sliding window and samples 45ms to 95ms in the case of 50ms sliding window as shown in Figure 1C.

### Estimation of time scales using conventional univariate analysis

One of the most popular methods for the estimation of time scales in univariate/single-channel(time series) data has been the estimation of time scales using the autocorrelation function (Golesorkhi et al., 2021b; Zeraati et al., 2022). To compare our multivariate method of time scale estimation against a benchmark method, we implemented a standard time scale estimation which has been used in many earlier studies. First, the autocorrelation values for every single trial was calculated across delays from 0 to 150ms (75 samples). The 150ms delays was selected as our trials had a 150ms ISI, therefore the autocorrelation signal could be affected by the upcoming stimulus if longer delays were incorporated. The autocorrelation function is defined as:

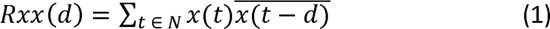

where x(n) is the signal time series, t is the time sample index, d is the delay between the original signal and its shifted version, N is the length of the time series analysed (i.e., here 75). The autocorrelation function was calculated for every channel and every trial. The autocorrelation function for normal neural signals should be an exponentially decaying function as a function of time delay as shown in Figure 1D. Importantly, depending on strength of the interrelationship of samples in the signal, the autocorrelation may have a strong or weak decay over time. Specifically, the autocorrelation function can quantify how far in time the samples of a time series are related to each other with interrelationships across further distances leading to more slowly decaying autocorrelation functions. Therefore, it can reflect the time scale of potential neural codes in EEG activation with longer neural codes showing a longer-lasting decay. In order to quantify the strength of the decay, we fitted an exponential decay function to the autocorrelation function in every trial and channel. The fitted exponential decay function is:

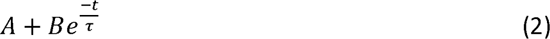

where A is the bias, B is the gain and r is the decay time constant. The time scale was estimated as the time delay at which the autocorrelation is reduced to 1/e = 0.368 times its initial value. Figure 1D shows the time scales and their corresponding autocorrelation curves in the range from 0 to 150ms.

### Bayes factor analysis

We used Bayes factor t-test analysis, as implemented by Bart Krekelberg based on Rouder et al. (2012), for comparing the levels of decoding across conditions and between a condition and baseline decoding. We used standard rules of thumb for interpreting levels of evidence (Lee and Wagenmakers, 2005; Dienes, 2014): Bayes factors of > 3 and < 1/3 were interpreted as evidence for the alternative and null hypotheses, respectively. We considered the Bayes factors which fell between 3 and 1/3 as suggesting insufficient evidence either way.

To evaluate the evidence for the null and alternative hypotheses of at-baseline and above-baseline decoding, respectively, we compared the decoding rates on every post-stimulus time point and the average of the decoding accuracies before the stimulus onset (−100 to 0ms; Figures 2 and 4). Accordingly, we performed the Bayes factor analysis for alternative (i.e., difference from baseline; H1) and the null (i.e., no difference from baseline; H0) hypotheses.

To evaluate the evidence for the null and alternative hypotheses of difference between decoding levels, we compared the decoding rates on every time point across the trial (−100 to 500 ms; Figure 2). We used a similar approach to compare the decoding levels across different time scales (Figure 5B) and across different conditions of each feature (Figure 5C).

The priors for all Bayes factor analyses were determined based on Jeffrey-Zellner-Siow priors (Jeffreys, 1998; Zellner and Siow, 1980) which are from the Cauchy distribution based on the effect size that is initially calculated in the algorithm using t-test (Rouder et al., 2012). The priors are data-driven and have been shown to be invariant with respect to linear transformations of measurement units (Rouder et al., 2012), which reduces the chance of being biased towards the null or alternative hypotheses. We did not perform correction for multiple comparisons when using Bayes factors as they are much more conservative than frequentist analysis in providing false claims with confidence (Gelman and Tuerlinckx, 2000; Gelman et al., 2012).

## Supplementary materials

**Supplementary Figure 1.**
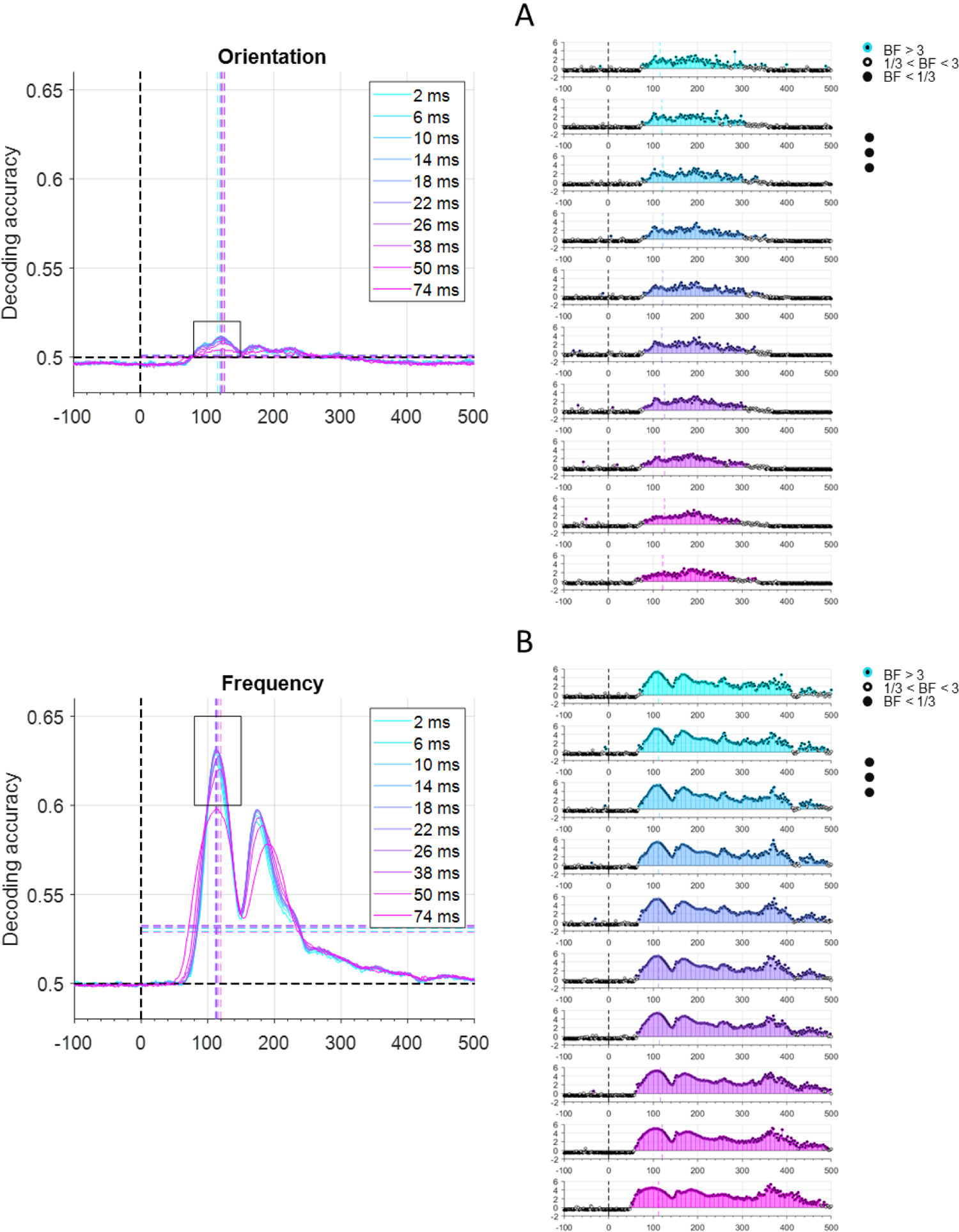
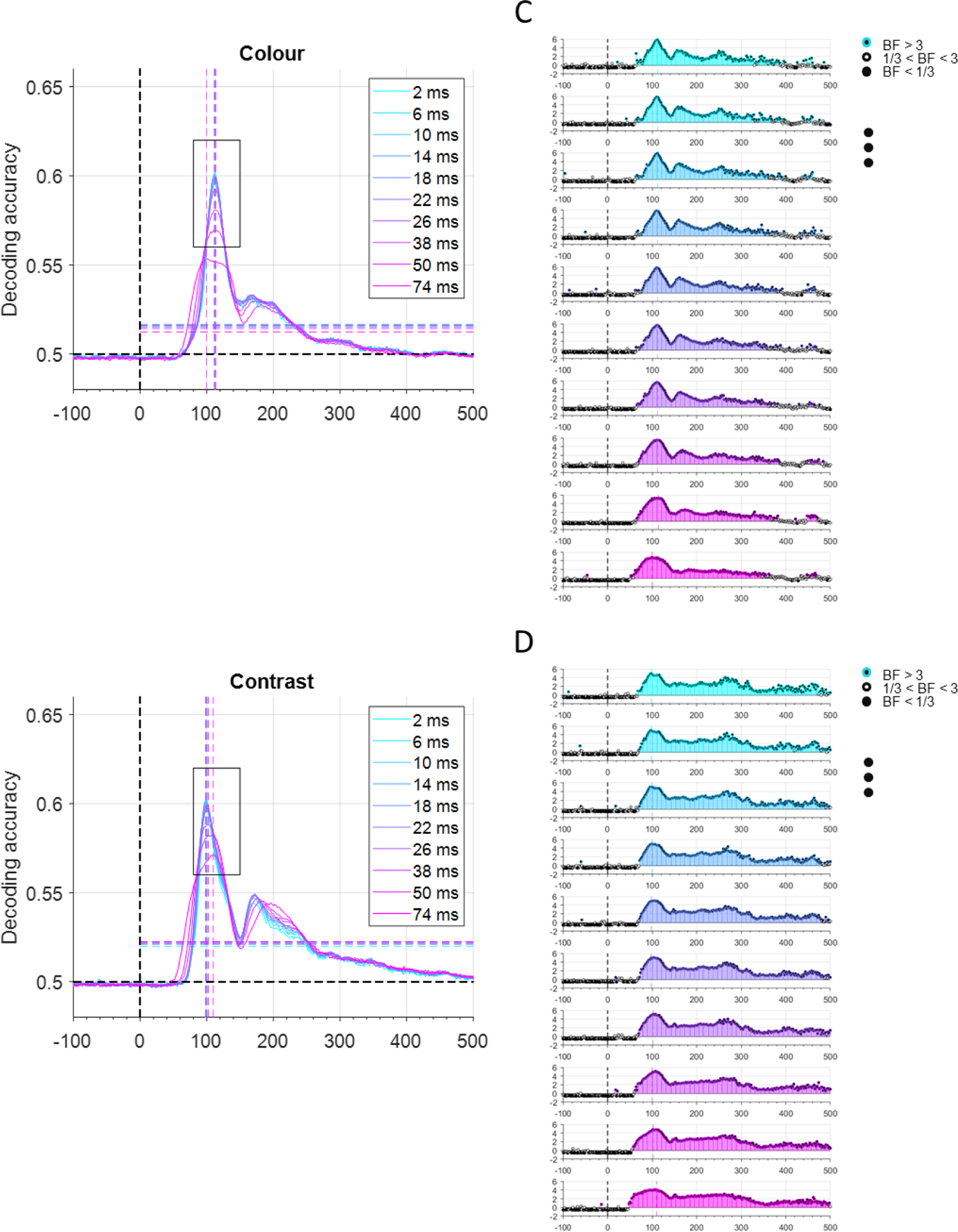
Decoding of stimulus features in across 10 time scales. Decoding time courses are shown for orientation (A), colour (B), frequency (C) and contrast (D). Left panels: decoding lines reflect average decoding across subjects. Black horizontal dashed line indicates theoretical chance-level decoding (0.5). Black vertical dashed line indicates stimulus onset time. The vertical and horizontal coloured dashed lines indicate the time of maximum decoding and the average post-stimulus decoding for the corresponding time scales (note that they may overlap), respectively. Bottom panels: Bayesian evidence for difference from baseline (−100 to 0ms). Bayes factors calculated using Bayes t-test (BF): Coloured filled circles show evidence for difference (BF > 3) or against any difference (BF < 1/3) and empty circles indicate insufficient evidence (1/3 < BF < 3).

**Supplementary Figure 2.**
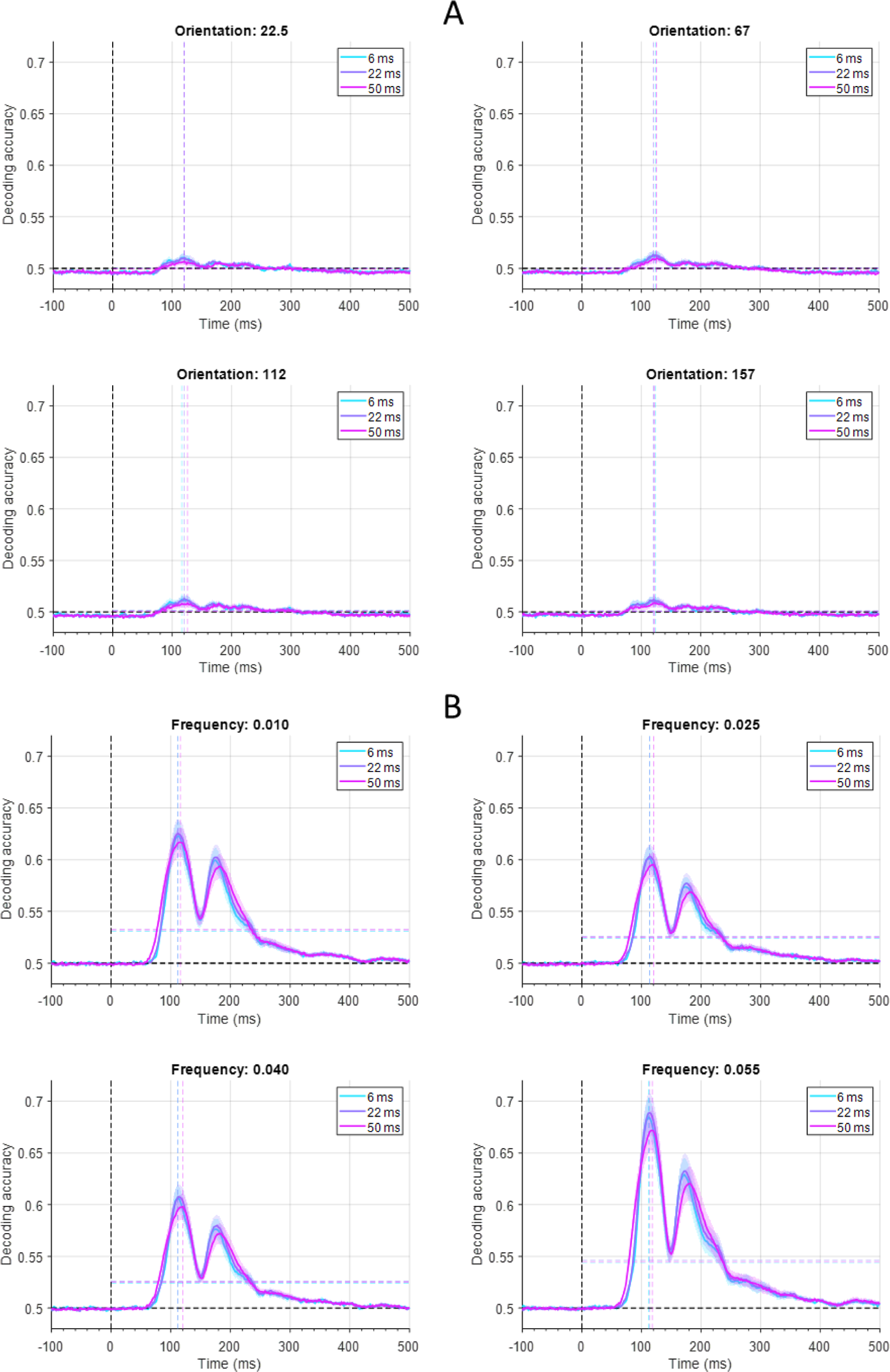
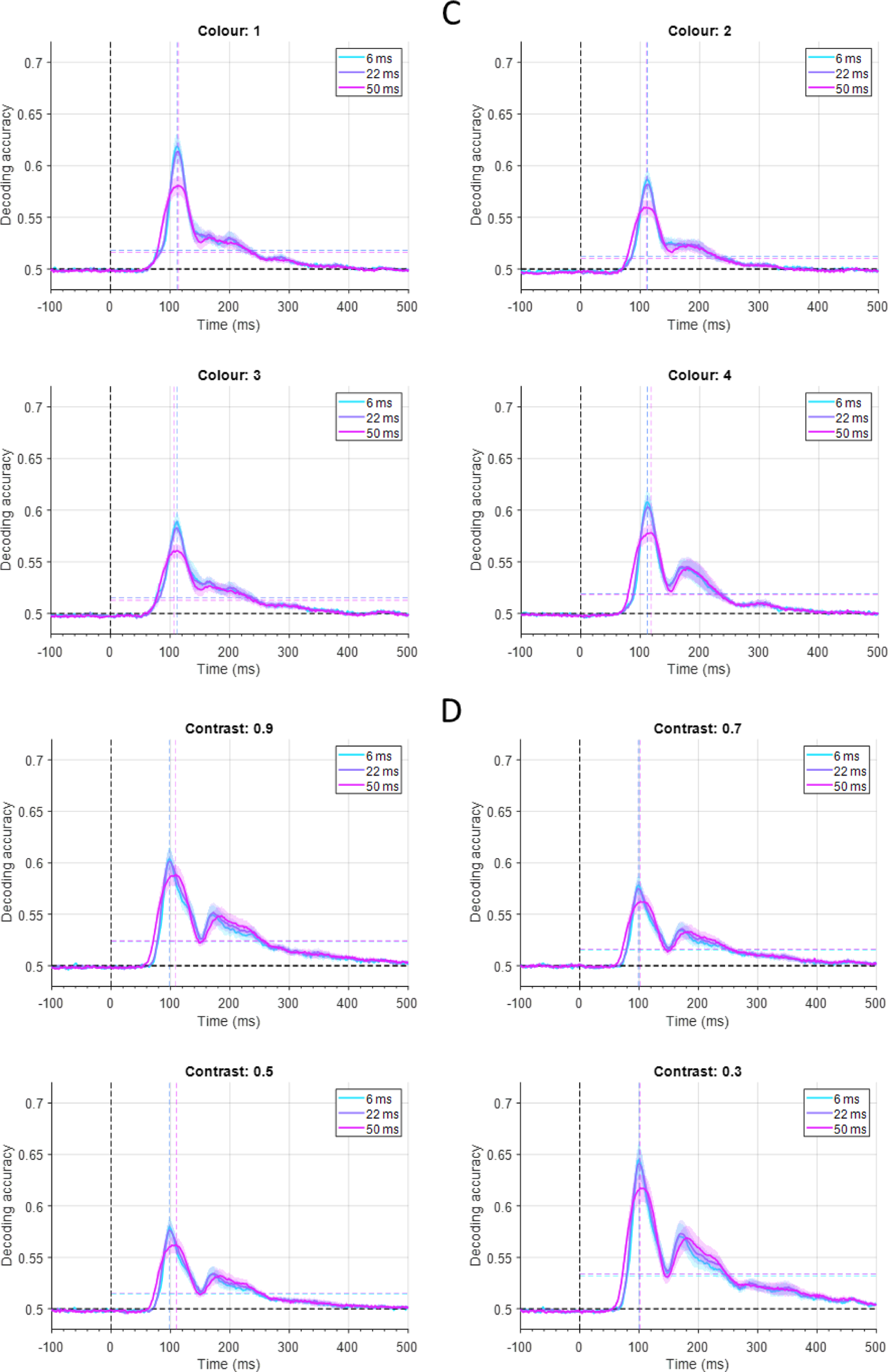
Decoding of stimulus features in three candidate time scales for every condition of each feature. Decoding time courses are shown for orientation (A), colour (B), frequency (C) and contrast (D). To obtain condition-specific decoding curves, among the 6 pairs of conditions which were decoded, only three decoding pairs which included the condition of interest were averaged. For example, to obtain decoding accuracy for the first orientation, decoding of first against second, first against third and fist against fourth conditions were averaged. Solid decoding lines and shadings reflect average and standard error across subjects, respectively. Black horizontal dashed line indicates theoretical chance-level decoding (0.5). Black vertical dashed line indicates stimulus onset time. The vertical and horizontal coloured dashed lines indicate the time of maximum decoding and the average post-stimulus decoding for the corresponding time scales (note that they may overlap), respectively.

**Supplementary Figure 3.**
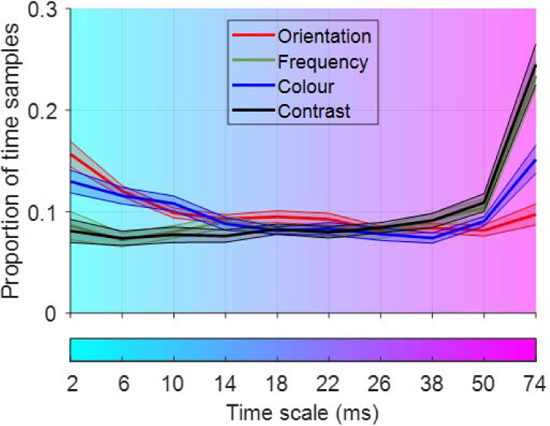
Proportion of time samples in which each time scale led to higher decoding than other time scales. The proportions were calculated based on the number of time points in the post-stimulus onset span in Figure 4. Solid lines and shadings reflect average and standard error across subjects, respectively. Time scales are shown using a spectrum of colours along the horizontal axis which match the colours in previous figures.

## Footnotes

1 https://klabhub.github.io/bayesFactor/

## References

Borst, A., & Theunissen, F. E. (1999). Information theory and neural coding. Nature neuroscience, 2(11), 947–957.

Butts, D. A., Weng, C., Jin, J., Yeh, C. I., Lesica, N. A., Alonso, J. M., & Stanley, G. B. (2007). Temporal precision in the neural code and the timescales of natural vision. Nature, 449(7158), 92–95.

Cavanagh, S. E., Hunt, L. T., & Kennerley, S. W. (2020). A diversity of intrinsic timescales underlie neural computations. Frontiers in Neural Circuits, 14, 615626.

Chase, S. M., & Young, E. D. (2007). First-spike latency information in single neurons increases when referenced to population onset. Proceedings of the National Academy of Sciences, 104(12), 5175–5180.

Delorme, A., & Makeig, S. (2004). EEGLAB: an open source toolbox for analysis of single-trial EEG dynamics including independent component analysis. Journal of neuroscience methods, 134(1), 9–21.

Denève, S., & Machens, C. K. (2016). Efficient codes and balanced networks. Nature neuroscience, 19(3), 375–382.

Dienes, Z. (2014). Using Bayes to get the most out of non-significant results. Frontiers in psychology, 5, 781.

Donoghue, T., Haller, M., Peterson, E. J., Varma, P., Sebastian, P., Gao, R., … & Voytek, B. (2020). Parameterizing neural power spectra into periodic and aperiodic components. Nature neuroscience, 23(12), 1655–1665.

Gawne, T. J., Kjaer, T. W., & Richmond, B. J. (1996). Latency: another potential code for feature binding in striate cortex. Journal of neurophysiology, 76(2), 1356–1360.

Gelman, A., & Tuerlinckx, F. (2000). Type S error rates for classical and Bayesian single and multiple comparison procedures. Computational statistics, 15(3), 373–390.

Gelman, A., Hill, J., & Yajima, M. (2012). Why we (usually) don’t have to worry about multiple comparisons. Journal of research on educational effectiveness, 5(2), 189–211.

Golesorkhi, M., Gomez-Pilar, J., Tumati, S., Fraser, M., & Northoff, G. (2021a). Temporal hierarchy of intrinsic neural timescales converges with spatial core-periphery organization. Communications biology, 4(1), 277.

Golesorkhi, M., Gomez-Pilar, J., Zilio, F., Berberian, N., Wolff, A., Yagoub, M. C., & Northoff, G. (2021b). The brain and its time: intrinsic neural timescales are key for input processing. Communications biology, 4(1), 970.

Grootswagers, T., Robinson, A. K., Shatek, S. M., & Carlson, T. (2023a). Features-EEG. OpenNeuro. [Dataset] doi: doi:10.18112/openneuro.ds004357.v1.0.0

Grootswagers, T., Robinson, A. K., Shatek, S. M., & Carlson, T. (2023b). Mapping the Dynamics of Visual Feature Coding: Insights into Perception and Integration. bioRxiv, 2023–04.

Grootswagers, T., Wardle, S. G., & Carlson, T. A. (2017). Decoding dynamic brain patterns from evoked responses: a tutorial on multivariate pattern analysis applied to time series neuroimaging data. Journal of cognitive neuroscience, 29(4), 677–697.

Harvey, M. A., Saal, H. P., Dammann III, J. F., & Bensmaia, S. J. (2013). Multiplexing stimulus information through rate and temporal codes in primate somatosensory cortex. PLoS biology, 11(5), e1001558.

Hasson, U., Chen, J., & Honey, C. J. (2015). Hierarchical process memory: memory as an integral component of information processing. Trends in cognitive sciences, 19(6), 304–313.

He, B. J. (2014). Scale-free brain activity: past, present, and future. Trends in cognitive sciences, 18(9), 480–487.

Hebart, M. N., & Baker, C. I. (2018). Deconstructing multivariate decoding for the study of brain function. Neuroimage, 180, 4–18.

Himberger, K. D., Chien, H. Y., & Honey, C. J. (2018). Principles of temporal processing across the cortical hierarchy. Neuroscience, 389, 161–174.

Jeffreys, H. (1998). The theory of probability. OuP Oxford.

Karimi-Rouzbahani, H. (2018). Three-stage processing of category and variation information by entangled interactive mechanisms of peri-occipital and peri-frontal cortices. Scientific reports, 8(1), 12213.

Karimi-Rouzbahani, H., & Woolgar, A. (2022). When the whole is less than the sum of its parts: maximum object category information and behavioral prediction in multiscale activation patterns. Frontiers in Neuroscience, 16, 825746.

Karimi-Rouzbahani, H., Bagheri, N., & Ebrahimpour, R. (2017a). Average activity, but not variability, is the dominant factor in the representation of object categories in the brain. Neuroscience, 346, 14–28.

Karimi-Rouzbahani, H., Bagheri, N., & Ebrahimpour, R. (2017b). Hard-wired feed-forward visual mechanisms of the brain compensate for affine variations in object recognition. Neuroscience, 349, 48–63.

Karimi-Rouzbahani, H., Ramezani, F., Woolgar, A., Rich, A., & Ghodrati, M. (2021a). Perceptual difficulty modulates the direction of information flow in familiar face recognition. NeuroImage, 233, 117896.

Karimi-Rouzbahani, H., Shahmohammadi, M., Vahab, E., Setayeshi, S., & Carlson, T. (2021b). Temporal variabilities provide additional category-related information in object category decoding: a systematic comparison of informative EEG features. Neural Computation, 33(11), 3027–3072.

Karimi-Rouzbahani, H., Vahab, E., Ebrahimpour, R., & Menhaj, M. B. (2019). Spatiotemporal analysis of category and target-related information processing in the brain during object detection. Behavioural brain research, 362, 224–239.

Kayser, C., Montemurro, M. A., Logothetis, N. K., & Panzeri, S. (2009). Spike-phase coding boosts and stabilizes information carried by spatial and temporal spike patterns. Neuron, 61(4), 597–608.

Lee, M. D., & Wagenmakers, E. J. (2005). Bayesian statistical inference in psychology: comment on Trafimow (2003).

Li, S., & Wang, X. J. (2022). Hierarchical timescales in the neocortex: Mathematical mechanism and biological insights. Proceedings of the National Academy of Sciences, 119(6), e2110274119.

Luczak, A., McNaughton, B. L., & Harris, K. D. (2015). Packet-based communication in the cortex. Nature Reviews Neuroscience, 16(12), 745–755.

Mainen, Z. F., & Sejnowski, T. J. (1995). Reliability of spike timing in neocortical neurons. Science, 268(5216), 1503–1506.

Majima, K., Matsuo, T., Kawasaki, K., Kawai, K., Saito, N., Hasegawa, I., & Kamitani, Y. (2014). Decoding visual object categories from temporal correlations of ECoG signals. Neuroimage, 90, 74–83.

Mazer, J. A., Vinje, W. E., McDermott, J., Schiller, P. H., & Gallant, J. L. (2002). Spatial frequency and orientation tuning dynamics in area V1. Proceedings of the National Academy of Sciences, 99(3), 1645–1650.

Merrikhi, Y., Shams-Ahmar, M., Karimi-Rouzbahani, H., Clark, K., Ebrahimpour, R., & Noudoost, B. (2021). Dissociable contribution of extrastriate responses to representational enhancement of gaze targets. Journal of cognitive neuroscience, 33(10), 2167–2180.

Montemurro, M. A., Rasch, M. J., Murayama, Y., Logothetis, N. K., & Panzeri, S. (2008). Phase-of-firing coding of natural visual stimuli in primary visual cortex. Current biology, 18(5), 375–380.

Murray, J. D., Bernacchia, A., Freedman, D. J., Romo, R., Wallis, J. D., Cai, X., … & Wang, X. J. (2014). A hierarchy of intrinsic timescales across primate cortex. Nature neuroscience, 17(12), 1661–1663.

Ng, B. S. W., Logothetis, N. K., & Kayser, C. (2013). EEG phase patterns reflect the selectivity of neural firing. Cerebral Cortex, 23(2), 389–398.

Norman-Haignere, S. V., Long, L. K., Devinsky, O., Doyle, W., Irobunda, I., Merricks, E. M., … & Mesgarani, N. (2022). Multiscale temporal integration organizes hierarchical computation in human auditory cortex. Nature human behaviour, 6(3), 455–469.

Oram, M. W. (2010). Contrast induced changes in response latency depend on stimulus specificity. Journal of Physiology-Paris, 104(3-4), 167–175.

Orbán, G., Berkes, P., Fiser, J., & Lengyel, M. (2016). Neural variability and sampling-based probabilistic representations in the visual cortex. Neuron, 92(2), 530–543.

Panzeri, S., Brunel, N., Logothetis, N. K., & Kayser, C. (2010). Sensory neural codes using multiplexed temporal scales. Trends in neurosciences, 33(3), 111–120.

Panzeri, S., Macke, J. H., Gross, J., & Kayser, C. (2015). Neural population coding: combining insights from microscopic and mass signals. Trends in cognitive sciences, 19(3), 162–172.

Pinto, L., Tank, D. W., & Brody, C. D. (2022). Multiple timescales of sensory-evidence accumulation across the dorsal cortex. ELife, 11, e70263.

Richman, J. S., & Moorman, J. R. (2000). Physiological time-series analysis using approximate entropy and sample entropy. American journal of physiology-heart and circulatory physiology, 278(6), H2039–H2049.

Rossi-Pool, R., Zainos, A., Alvarez, M., Parra, S., Zizumbo, J., & Romo, R. (2021). Invariant timescale hierarchy across the cortical somatosensory network. Proceedings of the National Academy of Sciences, 118(3), e2021843118.

Rouder, J. N., Morey, R. D., Speckman, P. L., & Province, J. M. (2012). Default Bayes factors for ANOVA designs. Journal of mathematical psychology, 56(5), 356–374.

Rupp, K., Roos, M., Milsap, G., Caceres, C., Ratto, C., Chevillet, M., … & Wolmetz, M. (2017). Semantic attributes are encoded in human electrocorticographic signals during visual object recognition. NeuroImage, 148, 318–329.

Schaefer, A. T., Angelo, K., Spors, H., & Margrie, T. W. (2006). Neuronal oscillations enhance stimulus discrimination by ensuring action potential precision. PLoS biology, 4(6), e163.

Schroeder, C. E., & Lakatos, P. (2009). Low-frequency neuronal oscillations as instruments of sensory selection. Trends in neurosciences, 32(1), 9–18.

Soltani, A., Murray, J. D., Seo, H., & Lee, D. (2021). Timescales of cognition in the brain. Current Opinion in Behavioral Sciences, 41, 30–37.

Spitmaan, M., Seo, H., Lee, D., & Soltani, A. (2020). Multiple timescales of neural dynamics and integration of task-relevant signals across cortex. Proceedings of the National Academy of Sciences, 117(36), 22522–22531.

Szczepański, J., Amigó, J. M., Wajnryb, E., & Sánchez-Vives, M. V. (2003). Application of Lempel–Ziv complexity to the analysis of neural discharges. Network: Computation in Neural Systems, 14(2), 335.

Theunissen, F., & Miller, J. P. (1995). Temporal encoding in nervous systems: a rigorous definition. Journal of computational neuroscience, 2, 149–162.

Thorpe, S., Fize, D., & Marlot, C. (1996). Speed of processing in the human visual system. nature, 381(6582), 520–522.

Victor, J. D. (2000). How the brain uses time to represent and process visual information. Brain research, 886(1-2), 33–46.

Walker, K. M., Bizley, J. K., King, A. J., & Schnupp, J. W. (2011). Multiplexed and robust representations of sound features in auditory cortex. Journal of Neuroscience, 31(41), 14565–14576.

Waschke, L., Kloosterman, N. A., Obleser, J., & Garrett, D. D. (2021). Behavior needs neural variability. Neuron, 109(5), 751–766.

Wolff, A., Berberian, N., Golesorkhi, M., Gomez-Pilar, J., Zilio, F., & Northoff, G. (2022). Intrinsic neural timescales: temporal integration and segregation. Trends in cognitive sciences, 26(2), 159–173.

Zamarashkina, P., Popovkina, D. V., & Pasupathy, A. (2020). Timing of response onset and offset in macaque V4: stimulus and task dependence. Journal of neurophysiology, 123(6), 2311–2325.

Zellner, A., & Siow, A. (1980). Posterior odds ratios for selected regression hypotheses. Trabajos de estadística y de investigación operativa, 31, 585–603.

Zeraati, R., Engel, T. A., & Levina, A. (2022). A flexible Bayesian framework for unbiased estimation of timescales. Nature computational science, 2(3), 193–204.

Zeraati, R., Shi, Y. L., Steinmetz, N. A., Gieselmann, M. A., Thiele, A., Moore, T., … & Engel, T. A. (2023). Intrinsic timescales in the visual cortex change with selective attention and reflect spatial connectivity. Nature communications, 14(1), 1858.

